# Pathway mining in functional genomics: An integrative approach to delineate boolean relationships between Src and its targets

**DOI:** 10.1101/2020.01.25.919639

**Authors:** Mehran Piran, Neda Sepahi, Mehrdad Piran, Pedro L. Fernandes, Ali Ghanbariasad

**Author notes:** Corresponding author: Mr. Mehran Piran.

## Abstract

**Motivation:** There are important molecular information hidden in the ocean of big data could be achieved by recognizing true relationships between different molecules. Human mind is very limited to find all molecular connections. Therefore, we introduced an integrated data mining strategy to find all possible relationships between molecular components in a biological context. To demonstrate how this approach works, we applied it on proto-oncogene c-Src.

**Results:** Here we applied a data mining scheme on genomic, literature and signaling databases to obtain necessary biological information for pathway inference. Using R programming language, two large edgelists were constructed from KEGG and OmniPath signaling databases. Next, An R script was developed by which pathways were discovered by assembly of edge information in the constructed signaling networks. Then, valid pathways were distinguished from the invalid ones using molecular information in articles and genomic data analysis. Pathway inference was performed on predicted pathways starting with Src and ending with the DEGs whose expression were affected by c-Src overactivation. Moreover, some positive and negative feedback loops were proposed based on the gene expression results. In fact, this simple but practical flowchart will open new insights into interactions between cellular components and help biologists look for new possible molecular relationships that have not been reported neither in signaling databases nor as a signaling pathway.

## Introduction

Data mining tools like programming languages have become an urgent need in the era of big data. Many Biological databases are available for free for the users, but how researchers utilize these repositories depends largely on their computational techniques. Among these repositories, databases in NCBI are of great interest. Pubmed literature database and GEO (Gene Expression Omnibus) (1) and SRA (Sequence Read Archive) (2) genomic databases are among the popular ones.

Different statistical models and machine learning algorithms have been developed to construct a gene regulatory network (GRN) from different genomic and epigenomic data archived in genomic databases (3, 4). These complex signaling networks are yielded by assembly of simple pathways. As results, pathway mining would aim to not only discover pathways but also understand the relationships and cross-talks between cellular pathways.

One of the simplest mining approaches to configure a signaling pathway is using molecular information in the literature. However, many papers illustrate paradox results because of the possible technical biases and type of biological samples. Some researchers obtain information they need by analyzing genomic data. Many biologists obtain the required information from signaling databases. Few researchers try to support data they find using different sources of information. As a result, a deep insight into the type of biological context is needed once inferring the correct relationships between cellular components.

In this study we integrated information coming from literature and genomic data analysis for some sets of constructed pathways. The study center of attention was on proto-oncogene c-Src having been implicated in progression and metastatic behavior of different carcinoma and adenocarcinoma (5-8). This gene is engaged in a process called Epithelial to Mesenchymal Transition (EMT) in which epithelial cells lose their cell-cell junction and acquire motility (9). Moreover, it is a target of many anti-cancer drugs, so revealing the new mechanisms that trigger cells to go under EMT would help design more potent drugs for different malignant tumors (10). To this end, two time series transcriptomic datasets were obtained from different technologies, microarray and NGS, in which MCF10A normal human adherent breast cell lines were equipped with ER-Src system. These cells were treated with tamoxifen to witness overactivation of c-Src. Then, gene expression pattern in different time points were analyzed for these cell lines. We tried to find firstly the most affected genes in these cells, secondly all possible connections between Src and DEGs (Differentially Expressed Genes). To be more precise in the analyses, only common DEGs in two datasets were considered. Furthermore, edges information in KEGG and OmniPath databases was used to construct pathways from Src to DEGs and between DEGs themselves. Finally, expression results and information in the literature were utilized to select possible correct pathways. Fig 1 illustrates the summary of whole experimental procedure.

**Fig 1:**
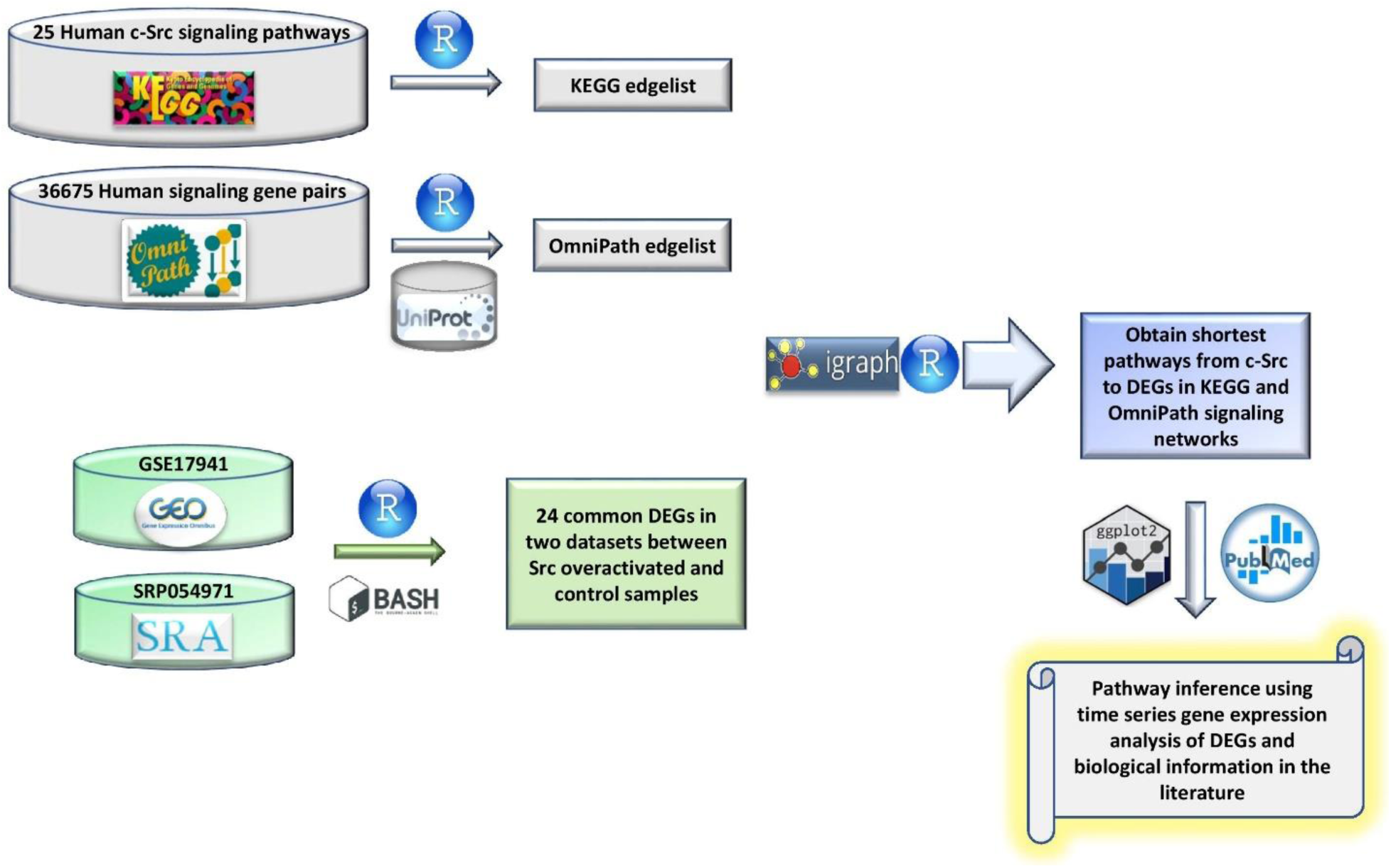
Visualization of different datamining steps using genomic and signaling databases to reach signaling pathways. KEGG and OmniPath edgelists were constructed from human KEGG signaling networks containing proto-oncogene c-Src and all human UniProt gene pairs (protein-protein) archived in OmniPath database. Two time series gene expression datasets were obtained from GEO and SRA databases and common DEGs were identified in the two datasets between tamoxifen treated samples and ethanol treated samples. Next, an R script was developed to recognize all shortest pathways from Src to the DEGs that were highly affected by Src overactivation. Using time series analysis of DEGs and information about target DEGs in the PubMed, possible valid pathways were distinguished from the false pathways.

## Methods

### Database Searching and recognizing pertinent experiments

Gene Expression Omnibus (http://www.ncbi.nlm.nih.gov/geo) and Sequence Read Archive (https://www.ncbi.nlm.nih.gov/sra) databases were searched to detect experiments containing high-quality transcriptomic samples concordance to the study design. Homo sapiens, SRC, overexpression, and overactivation were the keywords used in the search. Microarray raw data with accession number GSE17941 was downloaded from GEO database. RNA-seq fastq files from SRP054971 experiment were downloaded from SRA database. In both studies MCF10A cell lines containing ER-Src system were treated either with tamoxifen to overactivate Src or ethanol as the control.

### Microarray Data Analysis

R programming language version 3.6 was used to import and analyze the data. The preprocessing step involving background correction and probe summarization was done using RMA method in “affy” package (11). Absent probesets were identified using “mas5calls” function in this package. If a probeset contained more than two absent values, that one was regarded as absent and removed from the expression matrix. Besides, outlier samples were identified and removed using PCA and hierarchical clustering approaches. Employing Quantile normalization method, data were normalized. Many to Many problem which is mapping multiple probesets to the same gene symbol, was resolved using nsFilter function in genefilter package (12). This function selects the probeset with the largest interquartile range (IQR) to be representative of other probesets mapping to the same gene symbol. After that, “limma” package was utilized to identify differentially expressed genes between tamoxifen treated and ethanol treated cells (13).

### RNA-seq Data Analysis

Seven samples of RNA-seq study useful for the aim of our research were selected. Their accession IDs were from SRX876039 to SRX876045 (14). Bash shell commands were used to reach from fastq files to the counted expression files. Quality control step was done on each sample separately using “fastqc” function in “FastQC” module (15). Next, “Trimmomatic” software was used to trim reads (16). 10 bases from the reads head were cut and bases with quality less than 30 were removed and reads with the length of larger than 36 base pair were kept. Then, trimmed files were aligned to the hg38 standard FASTA reference sequence using “HISAT2” software to create SAM files (17). SAM files were converted into BAM files using “Samtools” package (18). In the last step, BAM files and a GTF file containing human genome annotation were given to “featureCounts” program to create counted files (19). After that, files were imported into R software (v3.6) and all samples were attached together to construct an expression matrix with 59,412 variables and seven samples. Rows with sum values less than 7 were removed and RPKM method in edger R package was used to normalize the rest of the rows (20). In order to identify DEGs, “limma” R package was used (13). Finally, correlation-based hierarchical clustering was done using “factoextra” R package [14].

### Pathway Construction from Signaling Databases

25 human KEGG signaling networks containing Src element were downloaded from KEGG signaling database (21). Pathways were imported into R using “KEGGgraph” package (22) and using programming techniques, all the pathways were combined together. Loops were omitted and only directed inhibition and activation edges were selected, so a large KEGG edgelist was constructed. In addition, a very large edgelist containing all literature curated mammalian signaling pathways were constructed from OmniPath database in R (23). To do pathway discovery, a script was developed in R using igraph package (24) containing a functions with four arguments. The first argument accepted an edgelist, second argument was a vector of source genes, third argument was a vector of target genes and forth argument received a maximum length of pathways.

## Results

### Data Preprocessing and Identifying Differentially Expressed Genes

Outlier sample detection was conducted using PCA (eigenvector 1 (PC1) and eigenvector 2 (PC2)) and hierarchical clustering. Fig 2A illustrates the PCA plot for the samples in GSE17941 study. Sample GSM448818 in time point 36-hour, was far away from the other samples which might be an outlier sample. In the hierarchical clustering approach, Pearson correlation coefficients between samples were subtracted from one for measurement of the distances. Then, samples were plotted based on their Number-SD. To get this number for each sample, the average of whole distances was subtracted from average of distances in all samples, then results of these subtractions were divided by the standard deviation of distance averages (25). Sample GSM448818_36h with Number-SD less than negative two was regarded as the outlier and removed from the dataset (Fig 2B).

**Fig 2:**
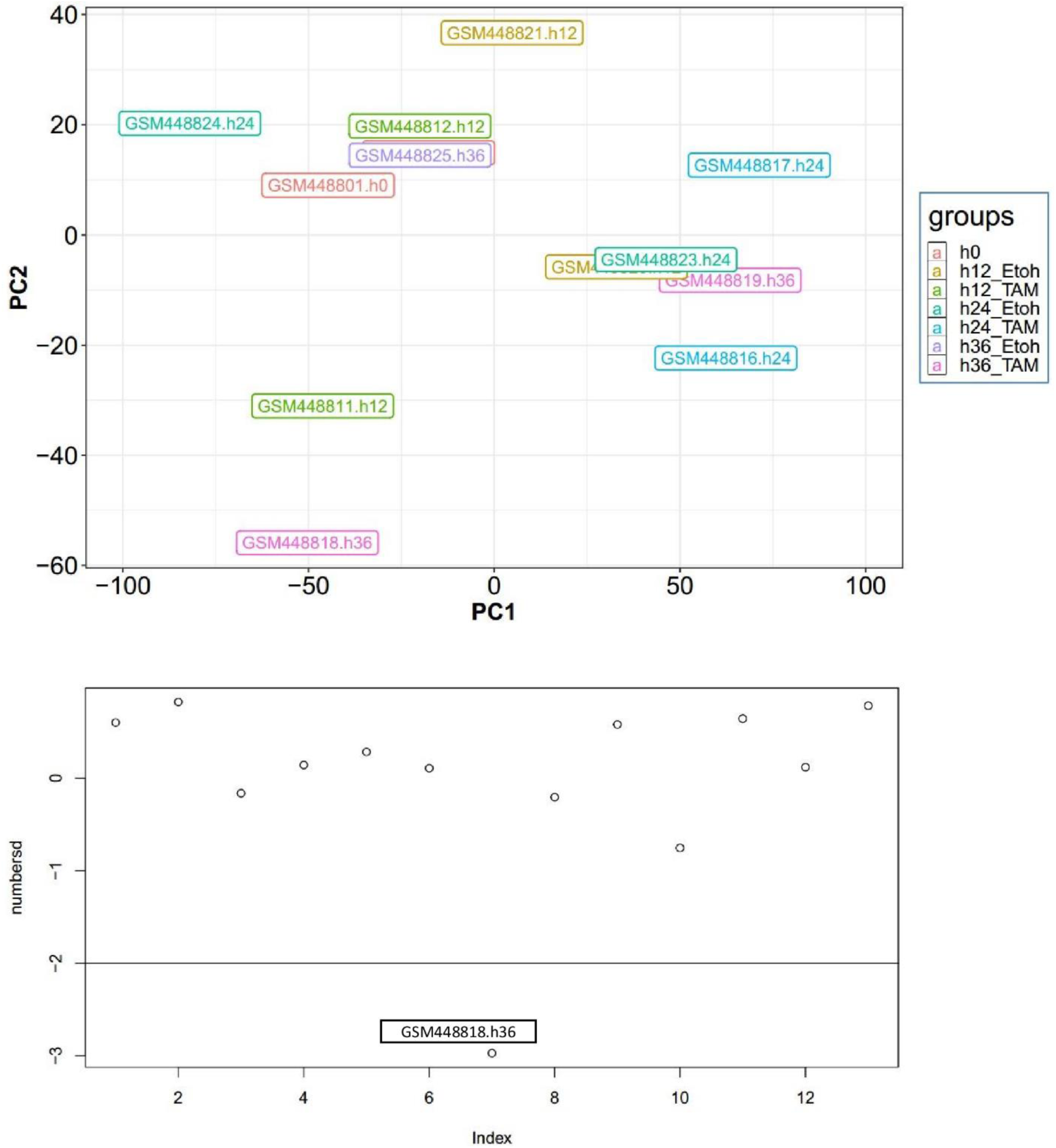
Outlier detection: A, is the PCA between samples in defferent time points in GSE17941 dataset. Replicates are in the same color. B, illustrates the numbersd value for each sample. Samples under -2 are regarded as outlier. The x axis represents the indices of samples.

There were 21 upregulated and 3 downregulated common DEGs between the two datasets. Fig 3 illustrates the average expression values between two groups of tamoxifen-treated samples and ethanol-treated samples for these common DEGs. For SRP054971 dataset average of 4-hours, 12-hour and 24-hour time points were used (A) and for GSE17941 dataset average of 12-hour and 24-hour time points were utilized (B). All DEGs had the absolute log fold change larger than 0.5 and p-value less than 0.05. Housekeeping genes were situated on the diagonal of the plot whilst all DEGs were located above or under the diagonal. This demonstrated that the preprocessed datasets were of sufficient quality for the analysis. In both datasets SERPINB3 was the most upregulated gene.

**Fig 3:**
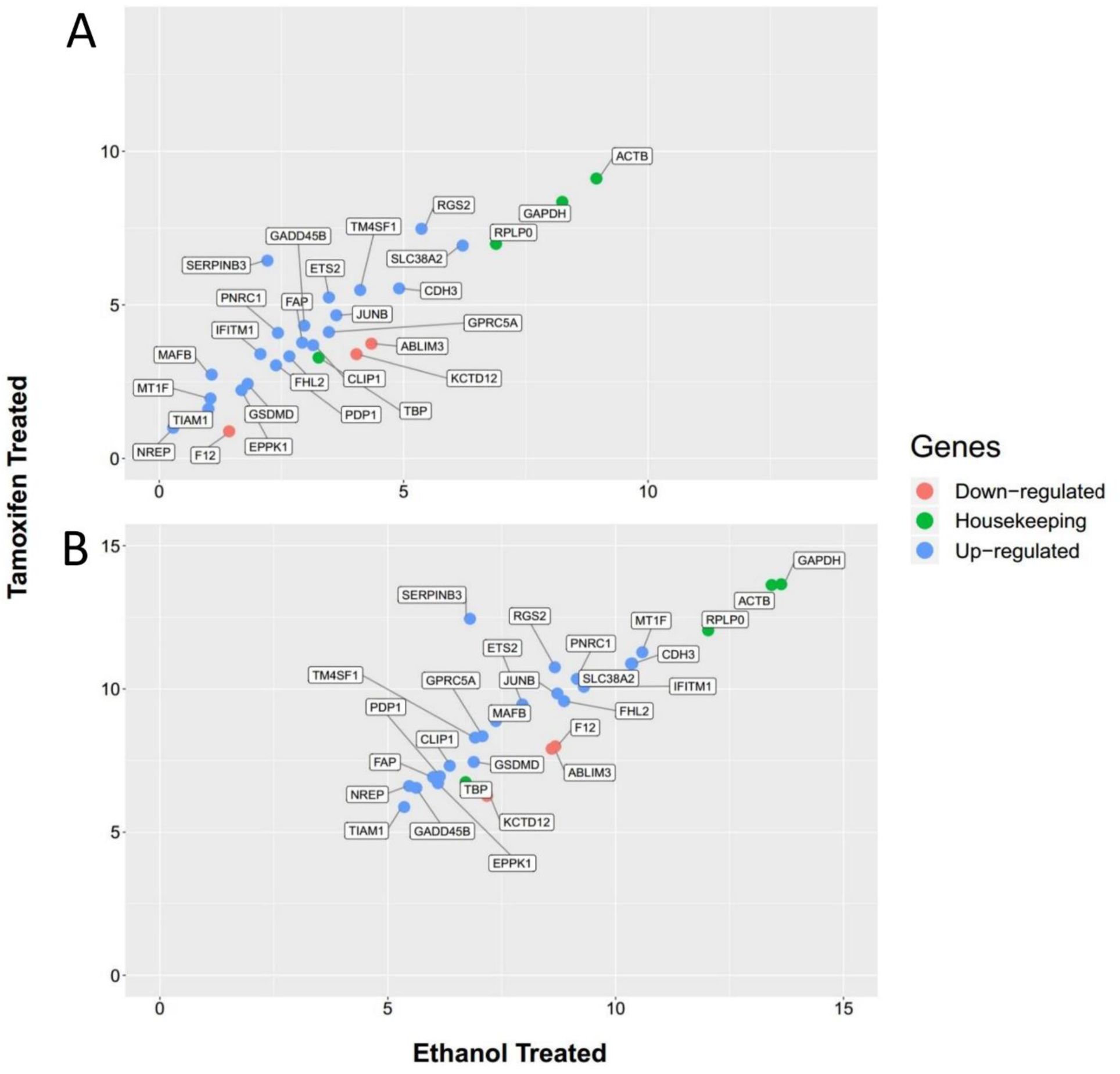
Scatter plot for upregulated, downregulated and housekeeping genes. The average values in different time points in A for SRP054971 dataset and B for GSE17941 dataset were plotted between control (ethanol treated) and tamoxifen treated.

### Pathway Analysis

Because obtained common DEGs were the results of analyzing datasets from two different genomic technologies, there was a high probability that Src-activation influence expression of these 24 genes. As a result, we continued our analysis in KEGG signaling network to find the boolean relationships between Src and these genes. For this purpose, we developed a pathway mining approach which extracts all possible directed pathways between two components in a signaling network. An R script was written to construct a large edgelist containing 1025 nodes and 7008 edges from 25 downloaded pathways containing Src (supplementary file 1). To reduce the number of pathways, only shortest pathways were regarded in the analysis. Just two DEGs were present in the constructed KEGG network namely TIAM1 and ABLIM3. Table1 shows all the pathways between Src and these two genes. They are two-edge distance Src targets in which their expression were affected by Src overactivation. TIAM1 was upregulated while ABLIM3 was downregulated in Fig 3. In Pathway number 1, Src induces CDC42 and CDC42 induces TIAM1 respectively. Therefore, a total positive interaction is yielded from Src to TIAM1. On the one hand ABLIM3 was downregulated in Fig 3. On other hand this gene could positively be induced by Src activation in four different ways in Table 1. We proposed that these pathways are not valid at mRNA level. Moreover, information in the signaling databases comes from different experimental sources and biological contexts. Consequently, a precise biological interpretation is required when combining information from different studies to predict a regulatory pathway.

**Table 1:** Discovered Pathways from Src gene to TIAM1 and ABLIM3 genes. In the interaction column, “1” presents activation state and “-1” shows inhibition state. ID column presents the edge IDs (indexes) in the KEGG edgelist. Pathway column represents the pathway number.

| Pathway | Source | Interaction | Target | Edge ID | Source | Interaction | Target | Edge ID |
| --- | --- | --- | --- | --- | --- | --- | --- | --- |
| 1 | SRC | 1 | CDC42 | E01596 | CDC42 | 1 | TIAM1 | E02917 |
| 2 | SRC | 1 | CDC42 | E01596 | CDC42 | 1 | ABLIM3 | E03143 |
| 3 | SRC | 1 | RAC3 | E03152 | RAC3 | 1 | ABLIM3 | E03136 |
| 4 | SRC | 1 | RAC2 | E03151 | RAC2 | 1 | ABLIM3 | E03135 |
| 5 | SRC | 1 | RAC1 | E03150 | RAC1 | 1 | ABLIM3 | E03134 |

Due to the uncertainty about the discovered pathways in KEGG, a huge human signaling edgelist was constructed from OmniPath database (http://omnipathdb.org/interactions) utilizing R programming. Constructed edgelist was composed of 20853 edges and 4783 nodes. Fourteen DEGs were found in the edgelist eleven of them were recognized to be Src targets. Eleven pathways with the maximum length of three and minimum length of one were discovered illustrated in Table2. Unfortunately, ABLIM1 gene was not found in the edgelist but Tiam1 was a direct (one-edge distance) target of Src in Pathway 1. FHL2 could be induced or suppressed by Src based on Pathways 3 and 4 respectively. However, FHL2 was among the upregulated genes by Src. Therefore, the necessity of biological interpretation of each edge is required to discover a possible pathway.

### Time Series Gene Expression Analysis

Src was overactivated in MCF10A (normal breast cancer cell line) cells using Tamoxifen treatment at the time points 0-hour, 1-hour, 4-hour and 24-hour in the RNA-seq dataset and 0-hour, 12-hour, 24-hour and 36-hour in the microarray study. The expression values for all upregulated genes in Src-activated samples were higher than controls in all time points in both datasets. The expression values for all downregulated genes in Src-activated samples were less than controls in all time points in both datasets. Fig 4 depicts the expression values for TIAM1, ABLIM3, RGS2, and SERPINB3 at these time points. RGS2 and SERPINB3 witnessed a significant expression growth in tamoxifen-treated cells at the two datasets. The time-course expression patterns of all DEGs are presented in supplementary file 2.

**Fig 4:**
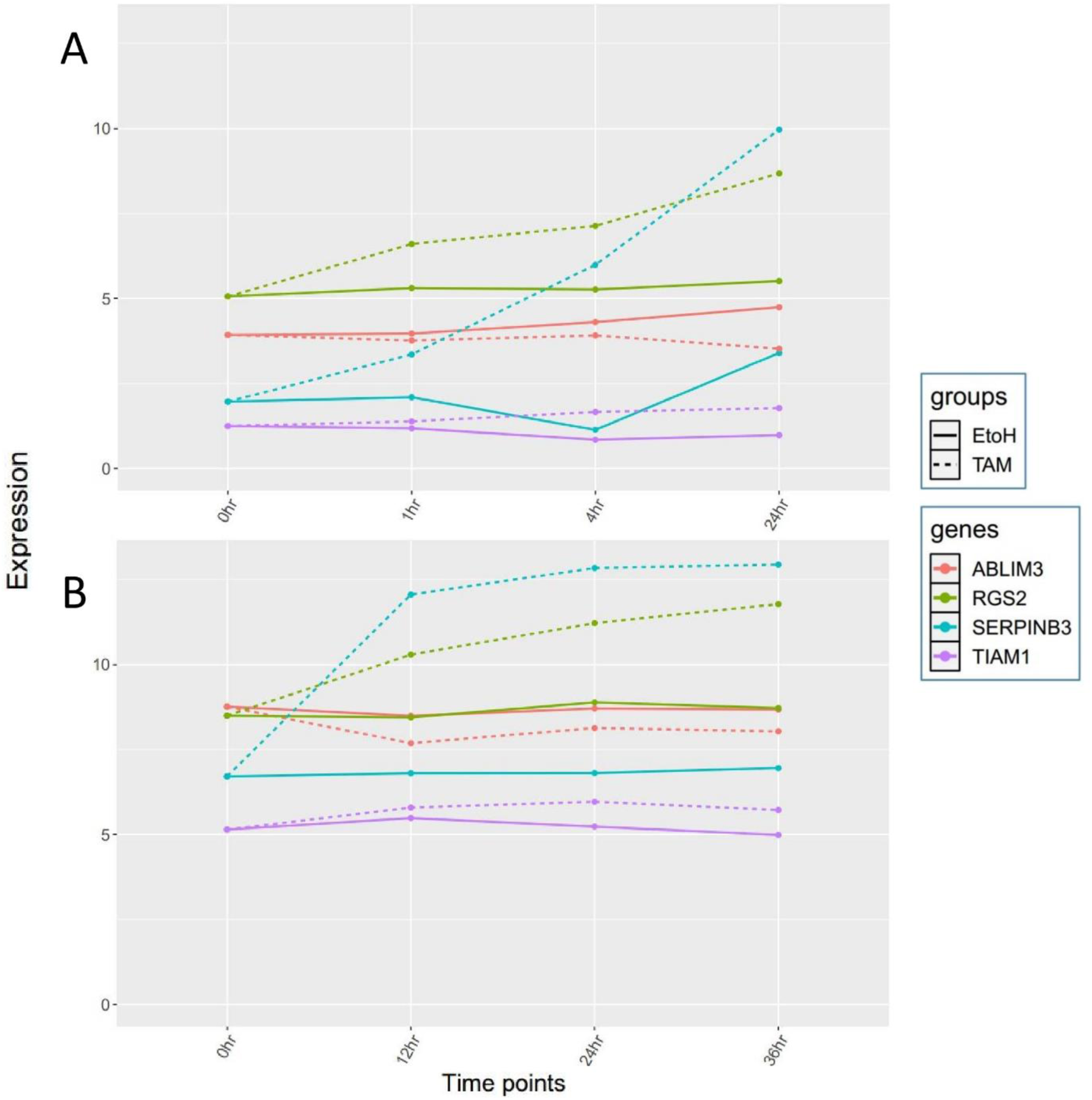
Expression values related to the four DEGs. TIAM1 and ABLIM3 are the genes present in the KEGG signaling network and RGS2 and SERPINB3 are the genes highly affected by activation of Src. A, presents values in SRP054971 dataset. B, presents values in GSE17941 dataset.

### Clustering

We applied hierarchical clustering on expression of all DEGs just in Src over-activated samples. Pearson correlation coefficient was used as the distance in the clustering method. Clustering results were different between the two datasets, therefore, we applied this method only on SRP054971 Dataset (Fig 5). We hypothesized that genes in close distances within each cluster may have relationships with each other. Among the four DEGs in Fig 4, Only TIAM1 and RGS2 were present in OmniPath edgelist. As a result, pathways were extracted from TIAM1 and RGS2 to their cluster counterparts. All these pathways are presented in supplementary file 3.

**Fig 5:**
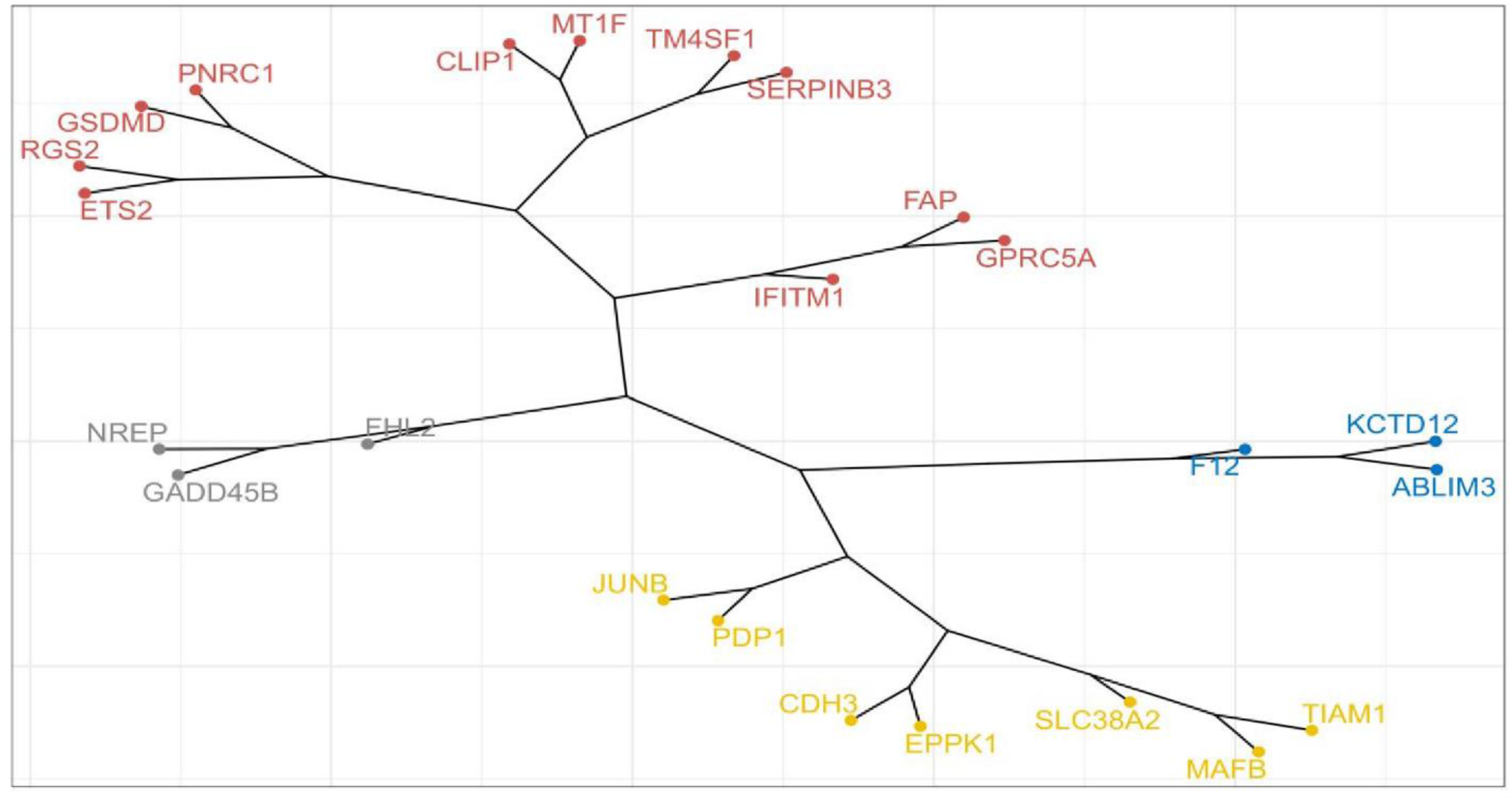
Pearson correlation-based hierarchical clustering. Figure shows the clustering results for expression of DEGs in RNA-seq dataset. Four clusters were emerged members of each are in the same color.

## Discussion

In many pathway inference methods, the necessity of valid data is required. Our approach is simple, is not based on mathematical models but real evidence. This pathway inference flowchart would demonstrate the importance of post transcriptional data and might also help distinguish the components in a pathway affected at transcript level and those affected at post-transcriptional level. Therefore, if other types of data rather than only mRNA is considered, many new biological pathways would be predicted by connecting edge information in more signaling databases. For instance, proteomic and phospho-proteomic data will be needed for pathway inference in a Protein-Protein Interaction Network (PPIN) if transition of signals are mostly related to the phosphorylation processes (26-28). Methylation sites on both mRNA and protein also impact cell signaling (29). Other modifications such as acetylation (30-32), ubiquitylation (33), sumoylation (34) impact the activity of different signaling networks as well. Consequently, different layers of gene regulation should be implemented on each gene pair once inferring causal relationships between gene pairs.

Analyzing the relationships between Src and all of these 24 obtained DEGs were of too much biological information. Therefore, we concentrated on TIAM1 and ABLIM3 (Actin binding LIM protein family member 3) and two highly affected genes in tamoxifen-treated cells namely RGS2 and SERPINB3. More investigations should be done on the rest of the genes to see how and why all these genes are affected by c-Src. That’s important because numerous studies have reported the role of Src implication in promoting EMT (35-38).

TIAM is a guanine nucleotide exchange factor (GEF) that is phosphorylated in tyrosine residues in cells transfected with oncogenic Src (39). Therefore, it could be a direct target of Src which has not been reported in 25 KEGG networks but this relationship was found in Table 2, pathway number 1. Src induces cell transformation through both Ras-ERK and Rho family of GTPases dependent pathways (40, 41). Moreover, The Rho GTPase Cdc42 can be phosphorylated and activated by Src through epidermal growth factor (EGF) signaling (42). CDC42 itself can promote activation of c-Src through EGFR signaling (43). Therefore, a positive feedback might occurs between the two molecules by Src activation. Not only CDC42 but also RAC1,2,3 GTPases are activated by Src through oncogenic growth factor receptors. These activations are transduced to GEF proteins such as TIAM1 which in turn regulate Rho-like GTPases GDP/GTP exchange keeping GTPases in their active form (44, 45). Therefore, by following the track of information in pathway number one in Tables 1 and 2, firstly a consistency in the flow of information is emerged secondly, the validity of pathways were demonstrated. Furthermore, Expression induction of TIAM1 in Fig 4 would explain how c-Src bolsters its cooperation with TIAM1 in order to keep Rho family of GTPases active leading to formation of membrane ruffles in vivo (39).

**Table 2:**
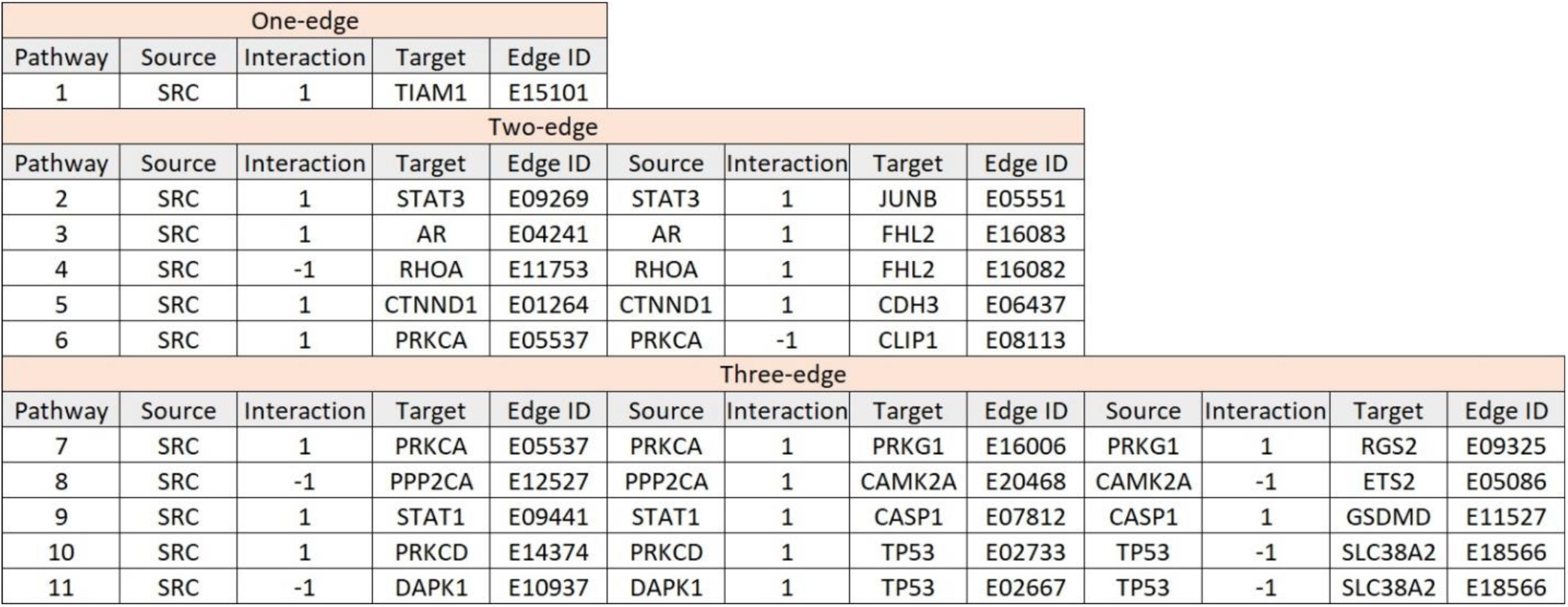
Discovered pathways from Src to the DEGs in OmniPath edgelist. In the interaction column, “1” depicts activation state and “-1” exhibits inhibition state. ID column presents the edge IDs (indexes) in the OmniPath edgelist. Pathway column represents the pathway number.

ABLIM3 is a component of adherent junctions (AJ) that interacts with actin filaments in epithelial cells and hepatocytes. Its downregulation in Fig 4 leads to the weakening of cell-cell junctions (46). Information flow for pathways targeting ABLIM3 is contrary to the results in Fig 4, therefore literature should be investigated carefully and probably post-transcriptional data is required to be analyzed, so we can predict that pathways numbers 2 to 5 in Table 1 are invalid during cell transformation in MCF10A cell lines.

SERPINB3 is a serine protease inhibitor that is overexpressed in epithelial tumors to inhibit apoptosis. This gene induces deregulation of cellular junctions by suppression of E-cadherin and enhancement of cytosolic B-catenin (47). In the Hypoxic environment of hepatic tumors, SERPINB3 is upregulated by HIF-2α and this upregulation requires intracellular generation of ROS (48). Moreover, there is a positive feedback between SERPINB3 and HIF-1α and -2α in liver cancer cells (49). Therefore, these positive mechanisms would help cancer cells to augment invasiveness properties and proliferation. Unfortunately, this gene did not exist neither in OmniPath nor in KEGG edgelists. Its significant expression induction and its effects in promoting metastasis would explain one of the mechanisms that Src triggers invasive behaviors in cancer cells.

RGS2 is a GTPase activating protein (GAP) for the alpha subunit of heterotrimeric G proteins (50). Although this gene was highly upregulated in Src-overexpressed cells, its expression has been found to be reduced in different cancers such as prostate and colorectal cancer (51, 52). This might overcomplicate the oncogenic effects of Src on promoting cancer. Pathway number 7 connects Src to RGS2 in Table 2. To infer pathway 7, ErbB2-mediated cancer cell invasion in breast cancer happens through direct interaction and activation of PRKCA/PKCα by Src (53). PKG1/PRKG1 is phosphorylated and activated by PKCα following phorbol 12-myristate 13-acetate (PMA) treatment (54). PRKG1 is a serine-threonine kinase activated through GMP binding. Inhibition of phosphoinositide (PI) hydrolysis in smooth muscles is done via phosphorylation of RGS2 by PRKG and its association with Gα-GTP subunit of G proteins. In fact, PRKG over-activates RGS2 to accelerate Gα-GTPase activity and enhance Gαβγ (complete G protein) trimer formation (55). Consequently, there would be a possibility that Src can induce RGS2 protein activity regarding the edge information in pathway 7. Src overactivation significantly induced expression of RGS2 in Fig 4. Therefore, there should be another pathway that increase the expression of RGS2 or maybe this upregulation is mediated by the mentioned components. Furthermore, RGS2 activates the GTPase activity of G proteins, so the upregulation of RGS2 by Src help to activate G proteins and promotion of cellular transformation.

SERPINB3 and ABLIM3 were not present in the OmniPath edgelist, so we conducted pathway mining from TIAM1 and RGS2 to their cluster counterparts and between TIAM1 and RGS2 (supplementary file 3). The results show that there would be a possible relationship between PNRC1, ETS2 and FATP toward RGS2. All these relationships were set by PRKCA and PRKG genes at the end of pathways. Nevertheless, PNCR1 made shorter pathways and is worth more investigation. JUNB and PDP1 make a relationship with TIAM1. The discovered pathway from JUNB to TIAM1 was mediated by EGFR and SRC demonstrating that SRC could induce its expression by induction of JUNB. Moreover, there were two pathways from TIAM1 to RGS2 and from RGS2 to TIAM1 with the same length which are worth more investigation.

In summary, Pathway mining is not just looking for direct pathways. Many hidden relationships will be unfolded once all findings and results in different databases around a specific component come together to reach a target component. Likewise we discovered possible new arrange of relationships between Src and Tiam1, between Src and RGS2 and between DEGs themselves.

## Supporting information

S1: KEGG networks and KEGG and OmniPath edgelists

S2: Time-series gene expression data

S3: discovered pathways in OmniPath signaling database

